# Inference of population structure from ancient DNA

**DOI:** 10.1101/261131

**Authors:** Tyler A. Joseph, Itsik Pe’er

**Affiliations:** Department of Computer Science, Columbia University, New York, NY 10027, USA; Department of Systems Biology, Columbia University, New York, NY 10027, USA; Data Science Institute, Columbia University, New York, NY 10027, USA

**Keywords:** population genetics, population structure, ancient DNA, time-series, variational inference, Kalman filtering

## Abstract

Methods for inferring population structure from genetic information traditionally assume samples are contemporary. Yet, the increasing availability of ancient DNA sequences begs revision of this paradigm. We present Dystruct (Dynamic Structure), a framework and toolbox for inference of shared ancestry from data that include ancient DNA. By explicitly modeling population history and genetic drift as a time-series, Dystruct more accurately and realistically discovers shared ancestry from ancient and contemporary samples. Formally, we use a normal approximation of drift, which allows a novel, efficient algorithm for optimizing model parameters using stochastic variational inference. We show that Dystruct outperforms the state of the art when individuals are sampled over time, as is common in ancient DNA datasets. We further demonstrate the utility of our method on a dataset of 92 ancient samples alongside 1941 modern ones genotyped at 222755 loci. Our model tends to present modern samples as the mixtures of ancestral populations they really are, rather than the artifactual converse of presenting ancestral samples as mixtures of contemporary groups.

**Availability:** Dystruct is implemented in C++, open-source, and available at https://github.com/tyjo/dystruct.

## 1 Introduction

The sequencing of the first ancient human genome [28], first Denisovan genome [29], and first Neanthertal genome [12] — all in 2010 — opened the floodgates for population genetic studies that include ancient DNA [19]. Ancient DNA grants a unique opportunity to investigate human evolutionary history, because it can provide direct evidence of historical relationships between populations around the world. Indeed, through combining ancient and modern samples, ancient DNA has driven many notable discoveries in human population genetics over the past ten years including the detection of introgression between anatomically modern humans and Neanderthals [24], evidence for the genetic origin of Native Americans [26], and evidence pushing the date of human divergence in Africa to over 250,000 years ago [30], among many others [2, 9, 19, 33].

Nonetheless, incorporating new types of DNA into conventional analysis pipelines requires careful examination of existing models and tools. Ancient DNA is a particularly challenging example: individuals are sampled from multiple time points from populations where allele frequencies have drifted over time. Hence, allele frequencies are correlated over time. The current state of the art for historical inference from ancient DNA uses pairwise summary statistics calculated from genome-wide data, called drift indices or F-statistics [21, 22], not to be confused with Wright’s F-statistics, that measure the amount of shared genetic drift between pairs of populations. Drift indices have several desirable theoretical properties, such as unbiased estimators, and can be used to conduct hypothesis tests of historical relationships and admixture between sampled populations [21]. Combined with tree-building approaches from phylogenetics, drift indices can reconstruct complex population phylogenies [18] including admixture events that are robust to difference in sample times. Computing drift indices, however, requires identifying populations *a priori*, a challenging task given that multiple regions around the world experienced substantial population turnover. Thus, exploratory tools that take an unsupervised approach to historical inference are required.

One of the most ubiquitous approaches to unsupervised ancestry inference is through the Pritchard-Stephens-Donnelly (PSD) model [23], implemented in the popular software programs *structure* and ADMIXTURE [1]. Under the PSD model, sampled individuals are modeled as mixtures of latent populations, where the genotype at each locus depends on the population of origin of that locus, and allele frequencies in the latent populations. Individuals can be clustered based on their mixture proportions, the proportion of sampled loci inherited from each population, which are interpreted as estimates of global ancestry [1]. ADMIXTURE computes maximum likelihood estimates of allele frequencies and ancestry proportions under the PSD model, while *structure* uses MCMC to compute posterior expectations of ancestry assignment. A key assumption of the PSD model is that populations are in Hardy-Weinberg equilibrium: the allele frequencies in each population are fixed. For ancient DNA, this assumption is clearly violated. The robustness of the PSD model to this violation remains under-explored.

In this paper, we develop a model-based method for inferring shared history between ancient and modern samples – Dystruct (Dynamic Structure) – by extending the PSD model to time-series data. To efficiently infer model parameters, we leverage the close connection between the PSD model and another model from natural language processing: latent Dirichlet allocation (LDA) [6]. The connection between the PSD model and LDA has long been known [3, 5], and applications of the statistical methodology surrounding LDA are beginning to enter the population genetics literature [11, 27]. Similar to the PSD model, LDA models documents as mixtures of latent topics, where each topic specifies a probability distribution over words. LDA has been successfully extended to a time-series model [5], where the word frequencies in topic distributions change over time in a process analogous to genetic drift. Thus, these dynamic topic models provide a natural starting point for models of population structure that incorporate genetic drift.

Our contributions are three-fold. First, we developed an efficient inference algorithm capable of parameter estimation under our time-series model. We extended the stochastic variational inference algorithm for the PSD model developed by [11] to time series data using the variational Kalman filtering technique developed by [5], and released software implementing our inference algorithm for general use. Second, we show that our model can lead to new insights on ancient DNA datasets: using simulations we demonstrate that Dystruct obtains more accurate ancestry estimates than ADMIXTURE on ancient DNA datasets; we then apply our model to a dataset of 92 ancient and 1941 modern samples genotyped at 222755 loci. Third, and more generally, our model opens the possibility for future model based approaches incorporating more complex demographic histories, complementing existing approaches for analyzing ancient DNA.

## 2 Methods

### 2.1 Preliminaries

Suppose we have genotypes of *D* individuals across *L* independent loci. Each individual *d* is a vector of *L* binomial observations, ***x*** = (*x_d_*_1_*, …, x_dL_*) for *x_dl_* ∈ {0, 1, 2}, where *x_dl_* is the number of non-reference alleles at locus *l*. Each individual is assumed to have been alive during one of a finite set of time points *g*[1]*, g*[2], …, *g*[*T*]. *g*[*t*] is measured as number of generations since the earliest time of interest. Each individual *d* is time stamped by *t_d_* ∈ {1, 2, …, *T*}, where *g*[*t_d_*] gives the time in generations when individual *d* was alive. We further define *Δg*[*t*] = *g*[*t*] *− g*[*t −* 1], the time in generations between time point *t* and time point *t −* 1.

Under the PSD model, each individual is a mixture from *K* latent populations. Let ***θ**_d_* = (*θ_d_*_1_*, …, θ_dK_*) be the ancestry porportions for individual *d*: ***θ**_d_* is the vector of probabilities that a locus in individual *d* originated in population *k*. Thus, *Σ_k_ θ_dk_* = 1. Let *β_kl_*[*t*] be the frequency of non-reference allele *l* in population *k* at time point *t*. The generative model for genotypes in each individual is:

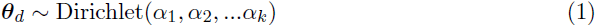

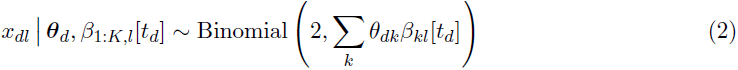

This follows the recharacterization of the original PSD model by [1] and [11]. In words, to generate an individual, we first draw mixture proportions ***θ**_d_* from a Dirichlet distribution with prior parameter ***α***. Then we draw genotypes from a binomial distribution, where the probability is a linear combination of the allele frequencies in the latent populations at time point *t_d_* (see Fig. 1).

To extend the model to time series data, we allow the allele frequencies to change at each time point using a normal approximation to genetic drift [7]:

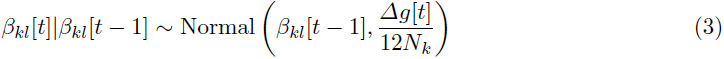

*N_k_* is the effective population size in population *k*. Initial allele frequencies *β_kl_*[0] and *N_k_* are parameters of the model. Initial allele frequencies *β_kl_*[0] are estimated from data, while *N_k_* are treated as known and fixed.

The state space model here is slightly different than normal approximation to the Wright-Fisher model for genetic drift. Under the Wright-Fisher model, changes in allele frequencies form a Markov Chain where alleles in future generations are chosen by sampling individuals from the previous generation with replacement. The total number of individuals with a particular allele in a diploid population, say *A*, one generation in the future is a Binomial(2*N_k_, p*) random variable, where *N_k_* is the population size, and *p* is the fraction of *A* alleles in the previous generation. Modeling allele frequencies rather than counts, the variance in allele frequencies Δ*g*[*t*] generations in the future, given the current allele frequency *β_kl_*[*t –* 1], is 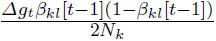. We approximate the variance by averaging over the interval (0, 1):

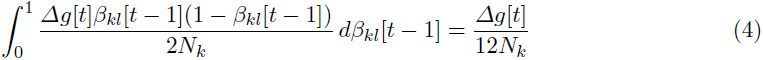

**Fig. 1.**
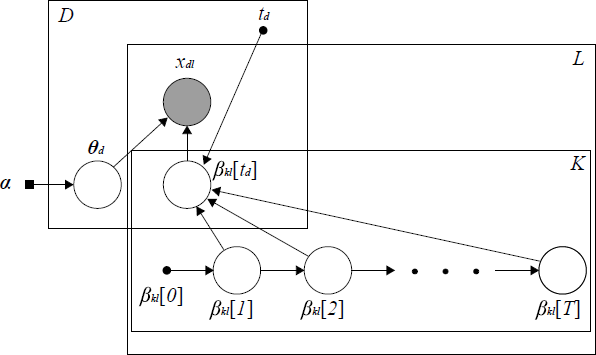
Graphical model depicting Dystruct’s generative model. *D* individuals are genotyped at *L* loci from *K* populations (boxes), and time stamped with time point *t_d_*. Each genotype in each individual, *x_dl_*, is a binomial observation that depends on: i) ancestry proportions, ***θ**_d_*, and; ii) allele frequencies *β_kl_*[*t_d_*] at time point *t_d_*. Allele frequencies *_kl_*[*t*] drift over time.

In practice, through simulations, we found that we were able to obtain accurate estimates despite this approximation. In this context, effective population size is not the parameter of interest, and we instead focus on ancestry estimates.

### 2.2 Posterior Inference

We take a Bayesian approach by inferring ancestry proportions through the posterior distribution *p*(***θ***_1:_*_D_*,***β***_1:_*_K_,*_1:_*_L_|**x***_1:_*_D,_*_1:_*_L_*). Direct posterior inference is intractable because the normal distribution is not a conjugate prior for the binomial. Following [5], we derive a variational inference algorithm that approximates the true posterior. We hereby summarize the variational inference approach for completion.

Variational inference methods [4,15,34] approximate the true posterior by specifying a computationally tractable family of approximate posterior distributions indexed by variational parameters, ***ϕ***. These parameters are then optimized to minimize the Kullback-Leibler (KL) divergence between the true posterior and its variational approximation. The key to variational inference algorithms relies on the observation that, given some distribution of latent parameters *q*(***z***), the log likelihood of the observations ***x*** can be decomposed into two terms:

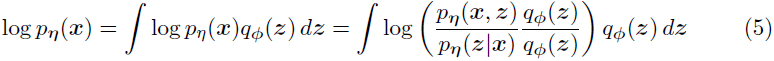

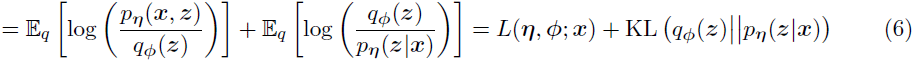

where ***η*** are the model parameters and ***ϕ*** are the variational parameters. The term on the right is the KL divergence between the true posterior and the variational approximation. Because KL divergence is non-negative, the *L* term in (6) is a lower bound on the log likelihood, called the evidence lower bound (ELBO). In practice, *L* is maximized with respect to the variational parameters ***ϕ***, minimizing the KL divergence between the true and approximate posterior. When maximized with respect to model parameters ***η***, approximate maximum likelihood estimates are obtained.

The posterior distribution in our model is given by

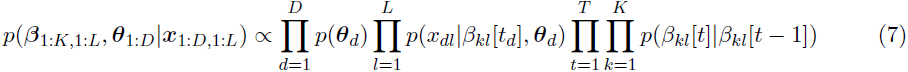

which we approximate by the variational posterior

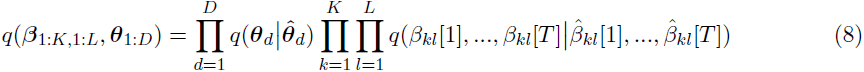

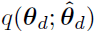 specifies a Dirichlet 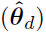 distribution. In the next section we elaborate on the form of 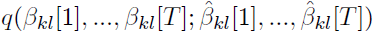.

### 2.3 Variational Kalman Filtering

Successful variational inference algorithms depend on formulating an approximate posterior close in form to the true posterior, and such that the expectations that make up the ELBO are tractable. Variational approximations commonly rely on the mean-field assumption, that posits the variational posterior as the product of independent distributions for each latent variable. However, this assumption is invalid for our model, as it ignores the dependence of allele frequencies over time. [5] introduced variational Kalman filtering as a technique to construct variational approximations to state space models with intractable posterior distributions. Variational Kalman filtering uses variational parameters 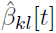 that are pseudo-observations from the state space model:

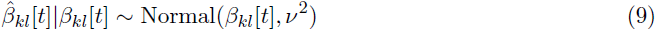

*ν* is an additional variational parameter. Given the pseudo-observations, standard Kalman filtering and smoothing equations can be used to calculate marginal means, 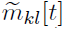, and marginal variances, 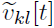, of the latent variables *β_kl_*[1: *T*] given the pseudo-observations 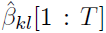. The variational approximation takes the form

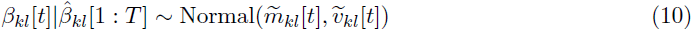

The ELBO is maximized with respect to the pseudo-observations.

### 2.4 Stochastic Variational Inference

Variational inference algorithms in the setting above often rely on optimizing parameters through coordinate ascent: each parameter is updated iteratively while the others remain fixed. Coordinate ascent can be computationally expensive, especially as the size of the data becomes large. For this reason, stochastic variational inference [4,14] is a popular alternative. Briefly, stochastic variational inference distinguishes global variational parameters, such as 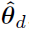, whose coordinate ascent update requires iterating through the entire dataset, with local parameters, 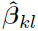, whose update only depends on a subset of the data.

We optimize variational parameters using stochastic variational inference (Algorithm 1 - see Appendix), along with a conjugate gradient algorithm for the local parameters 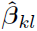 which do not have a closed form update, using a surrogate lower bound on the ELBO [11,35]. We first subsample a particular locus *l*, update the pseudo-outputs for that locus, then update the variational parameters 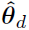 by taking a weighted average of the previous parameter estimates with the estimates obtained by looking at locus *l*. This process continues until the 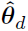 converge. See Appendix for details.

Estimates of ancestry proportions are computed by taking the posterior expectation of ***θ**_d_*: 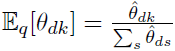

We further optimized our implementation, obtaining an order of magnitude speed up over a naive implementation. This improvement makes Dystruct feasible to use on realistic size datasets (see Section 3.3).

### 2.5 Simulated Data

We designed simulations to test the ability of our method to assign ancient samples into populations under two historical scenarios (Fig. 2). In each scenario, we simulated *K* populations at 10000 independent loci according to the Wright-Fisher model for genetic drift. We drew initial allele frequencies from a Uniform(0.2, 0.8) distribution, and simulated discrete generations by drawing 2*N_k_* individuals randomly with replacement from the previous generation. When then drew individuals at specific time points with genotypes and ancestry proportions specified by the generative model based on the allele frequencies at that time point. Note that we are not simulating data under the normal approximation.

**Fig. 2.**
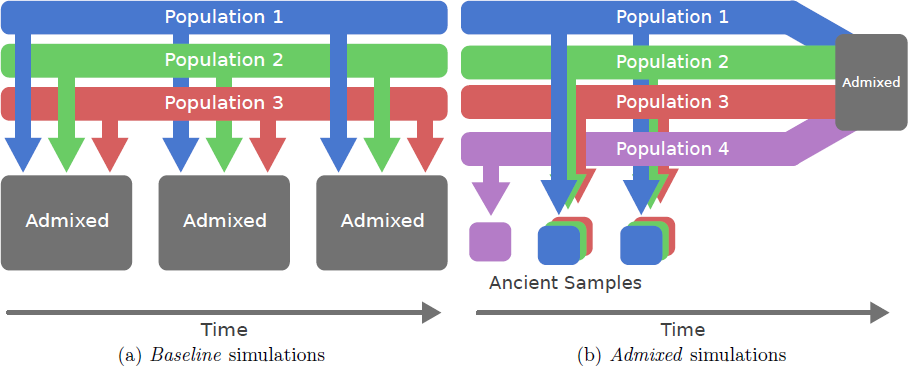
Simulation scenarios explored with Dystruct. (a) *Baseline* simulation scenario. Three populations were simulated that admixed at three time points. Individuals were sampled from the admixed populations. (b) *Admixed* simulation scenario. Ancient individuals were sampled from four unadmixed populations that merged to form a modern admixed population. Modern samples were from the admixed population.

One concern is that our model assumes allele frequencies are away from 0 or 1, while the allele frequencies in the Wright-Fisher model are guaranteed to fix given sufficient time. We allowed allele frequencies to fix in our simulations to test our model’s robustness to violating this assumption, though most allele frequencies do not reach fixation. We fixed effective population to *N_k_* = 2500 for all *k* = 1*, …, K*. To generalize our results across different effective population sizes, we measured time in coalescent units (1 coalescent unit = 2*N_k_* generations). We denote the total simulation time in coalescent units by *τ*. Each simulation was run across *τ* ∈ {0.02, 0.04, 0.08, 0.16}.

In the *baseline* simulation scenario (Fig. 2a), we sampled 40 individuals from *K* = 3 populations at 3 evenly spaced time points. We drew ancestry proportions from a Dirichlet(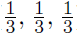) distribution, ensuring that the majority of any one individual’s genome originated in a single population, with smaller ancestry proportions from the remaining populations.

In the *admixed* scenario (Fig. 2b) we performed simulations that try to better mimic available real data. We assumed a modern population resulted from the instantaneous admixture of *K* = 4 ancestral populations. Ancient individuals were sampled pre-admixture and modern individuals were sampled post-admixture. We included two additional features found in current datasets. Such datasets comprise a small number of ancient samples when compared with modern samples. We therefore simulated 508 samples where 23 of the samples were ancient and the remaining 485 were modern, reflecting the ∼1:21 balance of samples in Section 2.6. All ancient samples occurred before time 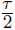. One of the four ancient populations was observed in the oldest ancient sample only, but appeared in modern populations, reecting the possibility that an ancient population may only be sampled once.

We repeated each simulation scenario 10 times for a total of 80 simulations, and compared the ability of our model to infer the parameter ***θ**_d_* with that of ADMIXTURE (v1.3.0). Since effective population size is a fixed parameter in Dystruct, we tested Dystruct on several effective population sizes. We ran Dystruct with *N_k_* = 1000; 2500; 5000; 10000 for all simulation scenarios. For each simulation, we computed the root-mean-square error (RMSE) between the true ances-try propq ortions, and parameters inferred by Dystruct and ADMIXTURE: RMSE ***θ**^true^, **θ**^inf^*) = 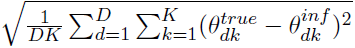

### 2.6 Real Data

[13] analyze a hybrid dataset of modern humans from the Human Origins dataset [17,21], 69 newly sequenced ancient Europeans, along with 25 previously published ancient samples [10,32], to study population turnover in Europe. Ancient samples included several Holocene hunter gatherers (∼5000-6000 years old), Neolithic farmers (∼5000-8000 years old), Copper/Bronze age individuals (∼3100-5300 years old), and an Iron Age individual (∼2900 years old). In addition, the data include three Pleistocene hunter-gatherers — ∼45000 year old Ust-Ishim [8], ∼30000 year old Kostenki14 [31], and ∼240000 year old MA1 [26] — the Tyrolean Iceman [16], the hunter-gatherers LaBrana1 [20] and Loschbour [17], and the Neolithic farmer Stuttgart [17].

We analyzed the publicly available dataset from https://reich.hms.harvard.edu/datasets. After removing related individuals identified in [13], and removing samples from outside the scope of their paper, we were left with a dataset consisting of 92 ancient samples and 1941 modern samples genotyped at 354212 loci. Again following [13], we pruned this original dataset for linkage disequilibrium in in PLINK [25] (v1.07) using-indep-pairwise 200 25 0.5, leaving 222755 SNPs. Each reported ancient sample includes confidence intervals for radiocarbon date estimates. To convert radiocarbon dates to generation time required by Dystruct, we assumed a 25 year generation time, and took the midpoint of the radiocarbon dates as point estimates divided by 25 for ancient samples. We further grouped time points for samples together if they were within the 95 % confidence interval for radiocarbon date estimates, and were part of the same culture. We assigned the year 2015 to modern samples. The final dataset spanned 1800 generations.

We then ran ADMIXTURE and Dystruct on the full data with effective population size of 7500 from *K* = 2 to *K* = 16 – the best supported *K* in [13] – and compared the results. Here we report the results for *K* = 11 because they have the clearest historical interpretation.

## 3 Results

### 3.1 Simulated Data

When simulating data according to the *baseline* scenario, Dystruct consistently matches up with ADMIXTURE or significantly outperforms it (Fig. 3a). When interpreting the order of magnitude of these accuracy results, it is important to note that ancestry vectors sum to 1, so a 0.01 decrease in RMSE improves relative accuracy of these vectors by order of *K*%. ADMIXTURE performs much worse as the simulated coalescent time increases, from RMSE of 0.032 for *τ* = 0.02 to RMSE of 0.082 at *τ* = 0.16. Dystruct is less susceptible to this increase in error. Intuitively, the more coalescent time is considered, the more the drift, and hence, the more important it is to model its dynamics.

**Fig. 3.**
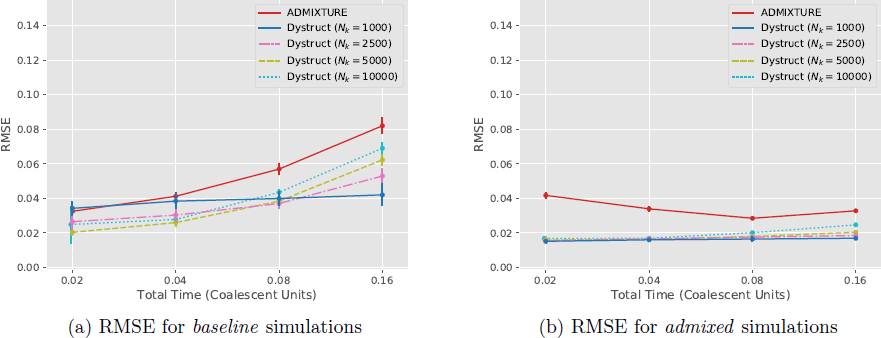
RMSE for the (a) *baseline* and (b) *admixed* simulation scenarios. Dystruct outperforms ADMIXTURE across several population size parameters for both scenarios. Ancestry vectors sum to 1, so a 0.01 improvement in RMSE corresponds to a 1% performance improvement.

The *admixed* simulation scenario demonstrates a substantial advantage to Dystruct on ancient samples across population parameters (Fig. 3b). This suggests that a promising use case for our model is investigating hypotheses of historical admixture. Nonetheless a near zero RMSE for Dystruct is potentially misleading because ancient samples are not admixed.

Dystruct also performs well on the modern samples. Dystruct outperforms ADMIXTURE for *τ* = 0.02, 0.04 by a factor of 2. At *τ* = 0.08, RMSE for ADMIXTURE and Dystruct are similar, while ADMIXTURE has a slight advantage at *τ* = 0.16.

### 3.2 Real Data

Dystruct shows good concordance with ADMIXTURE on modern data with known global populations (Fig. 5a). In particular, African populations (Dark Blue; eg. Bantu, Mbuti, Yoruba), Asian
populations (Red; e.g. Han, Japanese, Korean), Native American populations (Dark Pink; eg. Mixe, Mayan, Zapotec), and Oceanian populations (Yellow; eg. Papuan) all form similar genetic clusters, among many other examples.

**Fig. 4.**
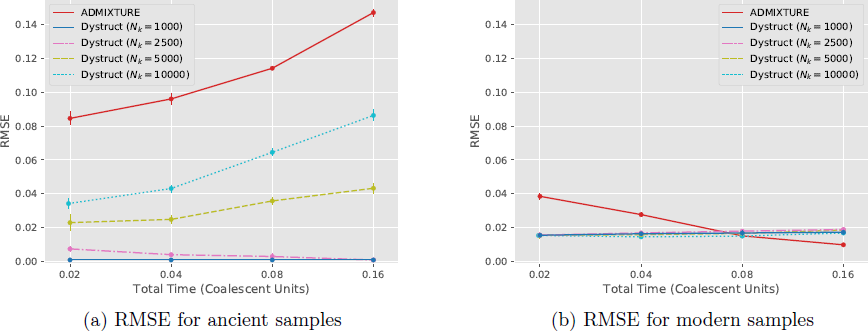
RMSE for ancestry estimates for (a) ancient samples and (b) modern samples for the *admixed* simulation scenario. Dystruct significantly outperforms ADMIXTURE when ancient samples are unadmixed (minimum RMSE = 0.00083). On modern samples, the error remains low for both Dystruct and ADMIXTURE.

Dystruct and ADMIXTURE differ on the ancient samples. In Dystruct, most ancient samples are “pure,” containing ancestry components from a single population, and modern day populations appear as mixtures of ancient populations. This is evident in the entropy across samples (Fig. 6). On ancient samples, Dystruct has lower entropy than ADMIXTURE, while the opposite is true for modern samples. This is most apparent in the different ancestry assignments for the oldest samples: the Pleistocene hunter gatherers. MA1, Kostenki14, and Ust-Ishim differ substantially in their representation between Dystruct and ADMIXTURE. These are the samples where genetic drift is most prominent. ADMIXTURE analysis describes MA1, Kostenki14, and Ust-Ishim as mixtures of several modern day populations. In contrast, Dystruct describes modern populations as mixtures of components derived from MA1, Kostenki14, and Ust-Ishim.

Most interestingly, the later ancient samples appear as mixtures of earlier samples in Dystruct, but not in ADMIXTURE. Late Neolithic, Bronze Age, and Iron Age samples appear as admixed between Yamnaya steppe herders (Orange), hunter-gatherers (Brown), and early Neotlithic (Green). Additionally, we see substantial shared ancestry between these groups and modern European populations. Both findings are consistent with [13], who found evidence supporting migration out of Yamnaya steppe herders into Eastern and Western Europe ∼4500 years ago, and supporting a model of European populations as a mixture of these groups. Kostenki14 shares ancestry with the Yamnaya group, suggesting a possible source for Yamnaya steppe ancestry.

### 3.3 Running Time

Despite the added complexity, additional model parameters, and large dataset, Dystruct ran on the real data in approximately 6 days using 2 cores of a 2.9 GHz Intel Core i5 processor. ADMIXTURE ran in approximately 2 days using 2 cores. Using 1 core, Dystruct ran in approximately 30 minutes per replicate on the *baseline* scenario, and approximately 120 minutes per replicate on the *admixed* scenario. ADMIXTURE ran in less than a minute on these scenarios. The advantages of stochastic variational inference are more apparent for larger datasets.

**Fig. 5.**
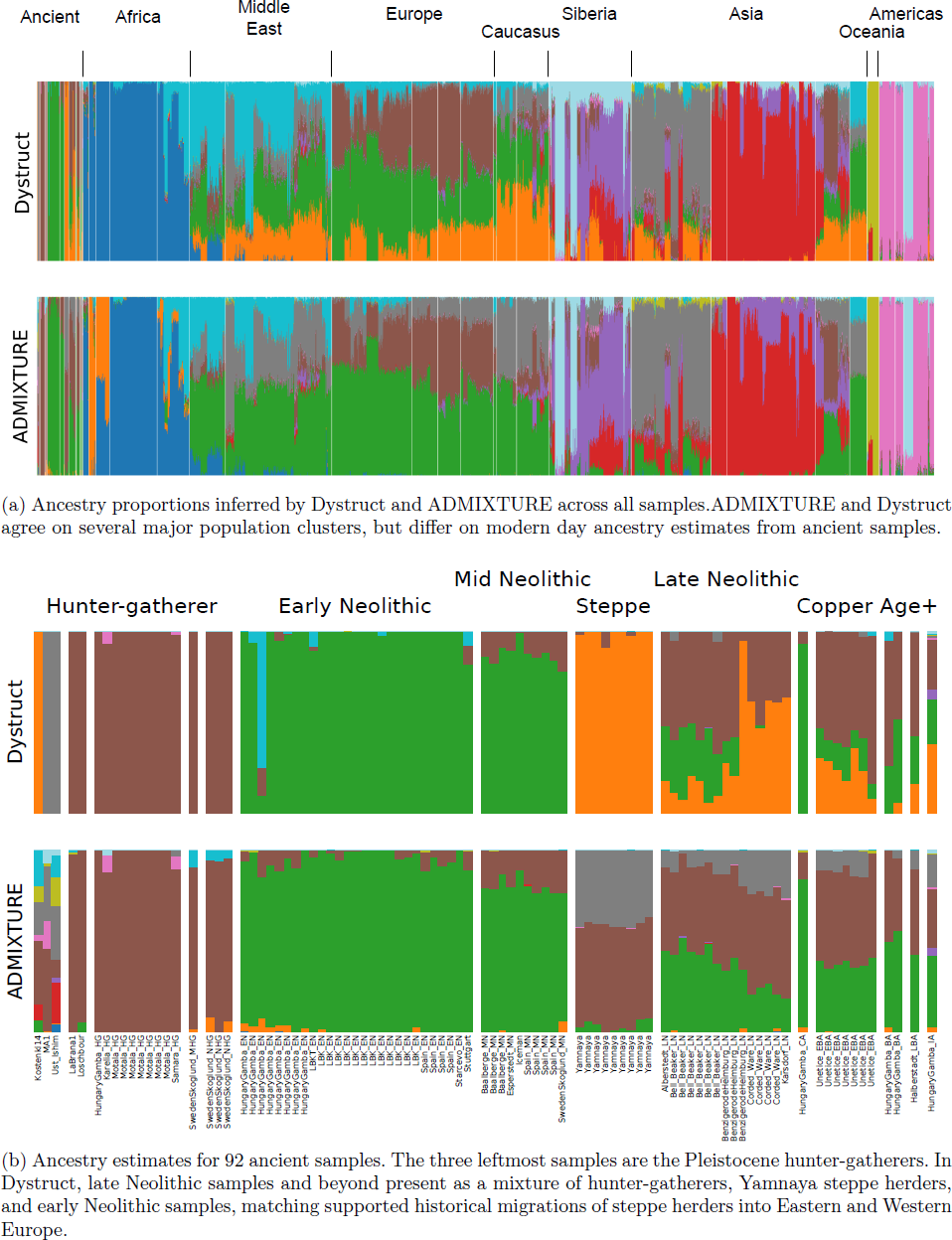
Ancestry proportions inferred across (a) all samples and (b) ancient samples only. Colors correspond between (a) and (b). Dystruct estimates ancestry for modern populations as combinations of ancient samples, while ADMIXTURE estimates ancestry for ancient samples as combinations of modern populations.

**Fig. 6.**
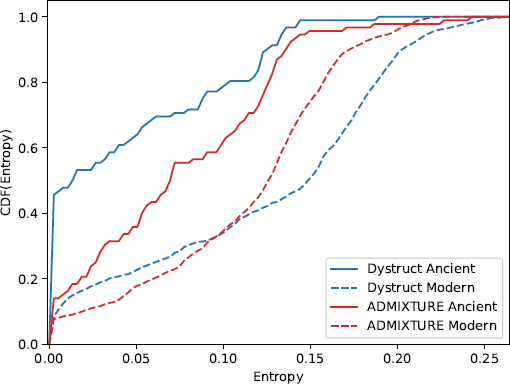
Cumulative density function for entropy across ancient and modern samples. Dystruct has a lower entropy for ancient samples, while ADMIXTURE has a lower entropy for modern samples.

Dystruct’s main computational consideration is the number of time points. During each iteration the parameters of a single locus are updated, then used to update ancestry estimates across all individuals. Estimates for ancestry parameters, 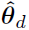, can be computed in closed form in *O*(*DK*); however, the update for the parameters 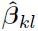 is approximated numerically. Computing the gradient of the 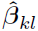 at a locus takes *O*(*T* ^2^ +*D*) time because the marginal means 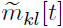 must be differentiated with respect to each pseudo-output. The gradient must be re-evaluated until the estimates for 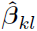 converge.

### 4 Discussion

We have presented Dystruct, a model and inference procedure to understand population structure and admixture from ancient DNA. The novelty of the model is its explicit temporal semantics. This formalization of allele frequency dynamics facilitates perception of modern and more recent populations as evolved from more ancient ones or combinations thereof. We derived an efficient inference algorithm for the model parameters using stochastic variational inference, and released software for use by the broader community. We established the performance of our model on several simulation scenarios, and further demonstrated its utility for gaining insight from the analysis of real data.

Our model outperforms the current standard modeling across a variety simulation scenarios. Encouragingly, our simulations show that Dystruct does a better job recovering population structure in the presence of genetic drift, an effect that hinders existing tools. Our model accurately detects when modern populations are mixtures of pure ancestral samples, while ADMIXTURE does not, and therefore is useful for testing hypotheses of historical admixture between ancient and modern populations.

We note the advantage of Dystruct increases with genetic drift and thus with coalescent time elapsed. This means that in practical situations, where samples are dated in years, Dystruct is most important when the effective population sizes are small. From statistical inference perspective, effective population size can be thought of as a regularizer that penalizes the difference between allele frequencies at each time point. Thus, as effective population size increases, alleles frequencies drift more slowly and become closer across time points, and estimates more closely match that of ADMIXTURE.

Our results on real data match known population clusters on modern populations, and lead to new interpretations of the ancient dataset. Interestingly, the PSD model tends to describe the oldest ancient samples as mixtures of modern populations, while in Dystruct several modern populations appeared as mixtures of these ancient samples. This makes sense in light of the standard goal of maximizing overall variance explained, a quantity dominated by the majority of the samples, which are modern. In contrast, temporal semantics implicitly assume admixture occurs forward in time, putting the focus on ancient populations. Dystruct can thus provide additional insight into such populations from ancient DNA.

There are several limitations to our approach. First, we model populations as independently evolving over time. This ignores historical relationships such as population splits. One potential side effect is that Dystruct may only capture one branch of a population phylogeny at a time. Second, across all simulations and for real data we constrained the effective population size across all populations to be the same. Thus, the parameters converge to one of at least *K* symmetric modes — population labels are exchangeable — and it is unclear how allowing different effective population sizes for different populations changes the log likelihood with respect to the parameter space. Future work should investigate this issue in more detail. Nonetheless, as we have demonstrated this is not a serious limitation to achieving reasonable estimates. Our results hold across a range of effective population sizes provided to Dystruct. Third, there is no clear procedure for choosing the correct number of populations *K*. We have deferred this issue to future work, but pose that this does not prevent a severe limitation: the current state of the art uses runs across multiple values of *K*, and interprets the results for each *K*.

More generally, we have presented a time-series model for population history with several promising extensions. Our method complements existing approaches, and can lead to new insights on ancient DNA datasets. Our work represents a first step toward statistical models capable of detecting complex population histories.

## 5 Acknowledgements

This material is based upon work supported by the National Science Foundation Graduate Research Fellowship under Grant No. DGE 16-44869, and the National Science Foundation under Grant No. DGE-1144854, and Grant No. CCF 1547120. Any opinion, findings, and conclusions or recommendations expressed in this material are those of the authors(s) and do not necessarily reflect the views of the National Science Foundation.

## A A Stochastic Variational Inference

In this section we derive the inference algorithm for stochastic variational inference under the Dystruct model. We first derive the traditional coordinate ascent updates, then show how we can modify these updates for stochastic optimization. Finally, we extend the algorithm for missing data.

### A.1 Computing The ELBO

The ELBO is given by

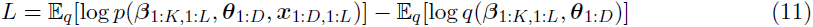

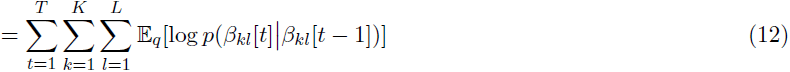

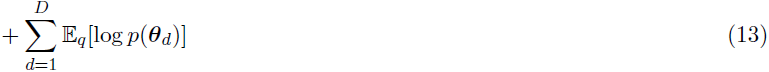

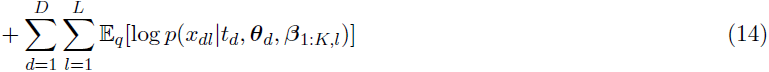

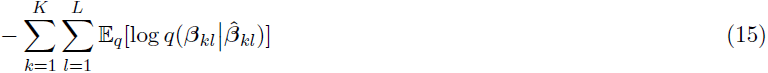

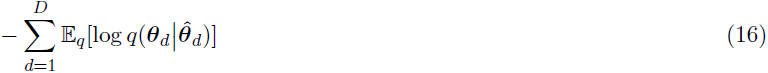

The ELBO as written does not have a closed form due to the log sum terms that appear in (14): 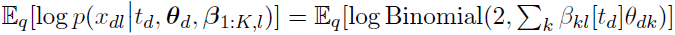:

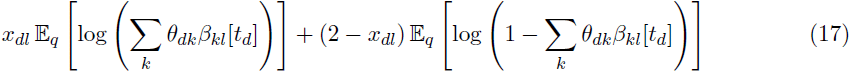

Following [11], we optimize a surrogate lower bound by introducing auxiliary variational parameters *ϕ_dl_* = (*ϕ_dl_*[1]*, …, ϕ_dl_*[*K*]) and ***ζ**_dl_* = (*ζ_dl_*[1]*, …, ζ_dl_*[*K*]) whose vector components sums to 1. An application of Jensen’s inequality shows

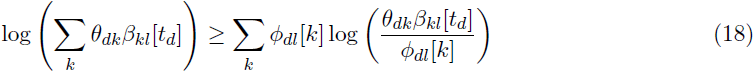

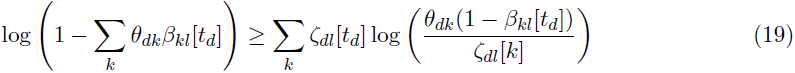

so we still maintain a lower bound on the log likelihood. The auxiliary parameters are optimized to provide a tight lower bound. Fixing all other parameters, the constrained optimization problem can be solved using an application of Lagrange multipliers

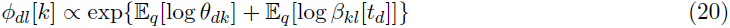

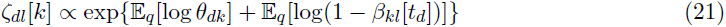

The first term in both equations is an expectation of a sufficient statistic, and therefore has a closed form: 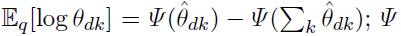 is the Digamma function. The two expectations in the second terms can be approximated by taking second order Taylor expansions around the marginal means of *β_kl_*[*t_d_*] and 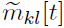:

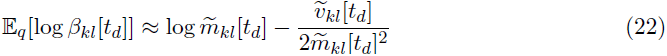

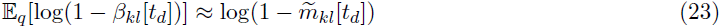

### A.2 Optimizing Ancestry Proportions

Note that the *q*(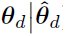) satisfy the mean field assumption - the ***θ**_d_* in the variational posterior are independent. Therefore they have optimal coordinate ascent updates of the form

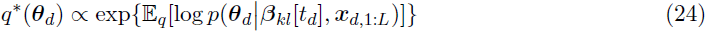

where we have used several conditional independencies to simplify the complete conditional of ***θ**_d_*. Using the surrogate lower bound in the ELBO gives the optimal update

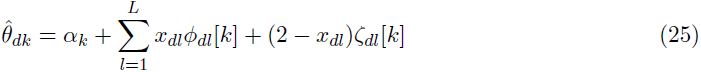

matching the expression in [11].

### A.3 Optimizing allele frequencies

In variational Kalman filtering, the variational distribution for each *β_kl_*[*t*] is given by

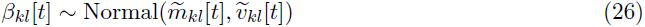

where the mean and variance are the marginal means and posteriors given by the Kalman filtering and smoothing equations. Following the notation in [5], the forward (filtered) means and variances are given by

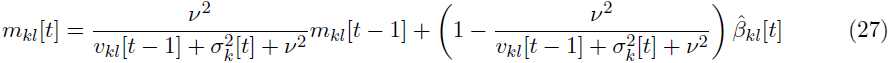

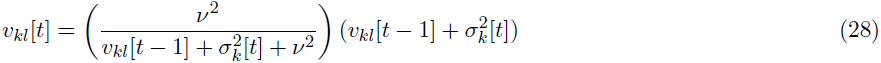

where 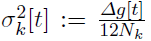. The initial conditions are *m_kl_*[0] = *β_kl_*[0]. The marginal (smoothed) means and variances are

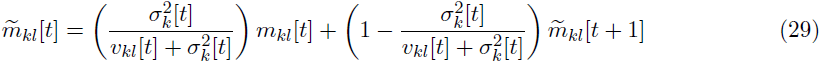

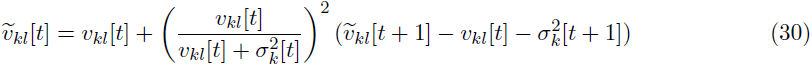

with initial conditions 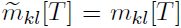 and 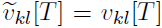. The variational parameters 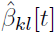 are optimized with respect to the ELBO, hence we need the partial derivatives of the marginal means 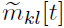 with respect to 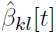. These can be obtained using the forward-backward recurrence as in [5]. We will show the recurrence for initial frequencies *β_kl_*[0], which are not maximized in [5], and note that the other partial derivations can be obtained similarly. The recurrence is

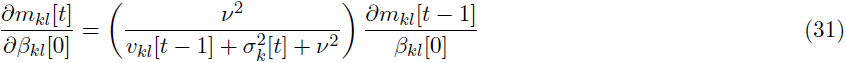

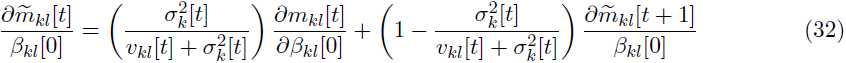

We optimize the 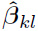 with respect to a single locus in a single population at time using a conjugate gradient algorithm, constraining the parameters to lie in the interval (0, 1). The terms in the ELBO with respect to locus *l* in population *k* are

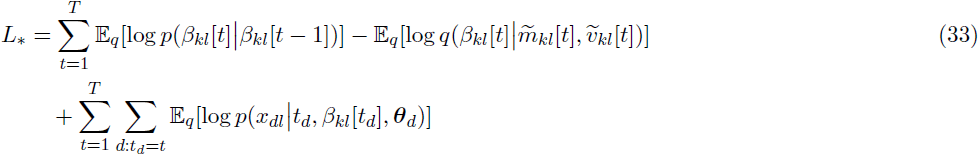

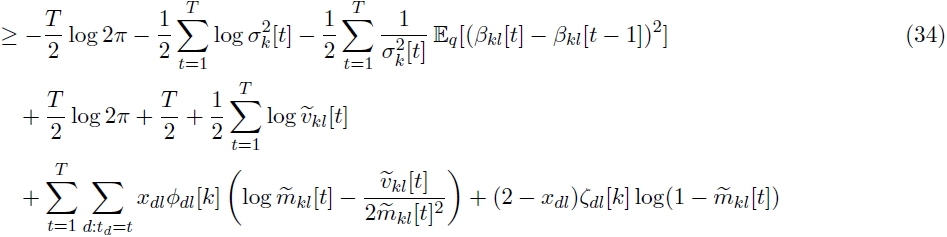

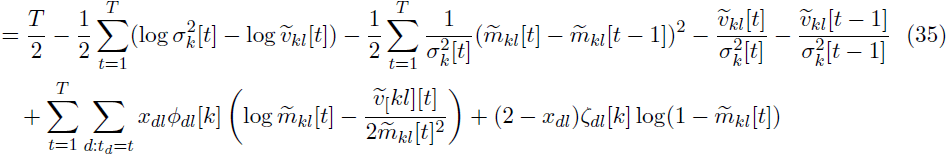

where we define 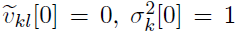, and 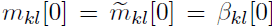 for notational convenience. Taking partial derivatives with respect to the pseudo-outputs gives us

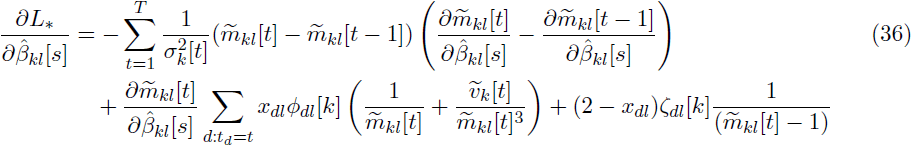

The full algorithm iterates between optimizing the local parameters 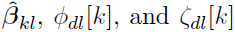 for each locus in each individual in each population using (20), (21), and (36), then updating global parameters 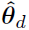 according to (25) until convergence.

### A.4 Inference Algorithm

We can perform stochastic variational inference through a slight modification to the coordinate ascent algorithm presented above [4, 14]. Stochastic variational inference computes noisy estimates of the optimal global parameters by stochastically subsampling data points, and using the optimal local parameters to update the global parameters. The optimal global parameters are a weighted average of the previous global parameters, with the newly computed global parameters. Following [11], the *n* + 1 stochastic variational inference update for the global parameters 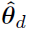 is

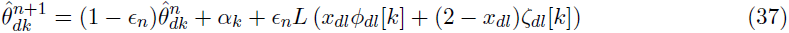

where *∊_n_* is the step size for iteration *n* and *L* is the number of loci. Provided the step size meets certain criteria the algorithm is guaranteed to converge. See [14] or [4] for more details. We picked a step size of *∊_n_* = (1 + *n*)^−0.5^ for the first 10000 iterations, and *∊_n_* = (*n* − 7825)^−0.6^ for the remaining iterations.

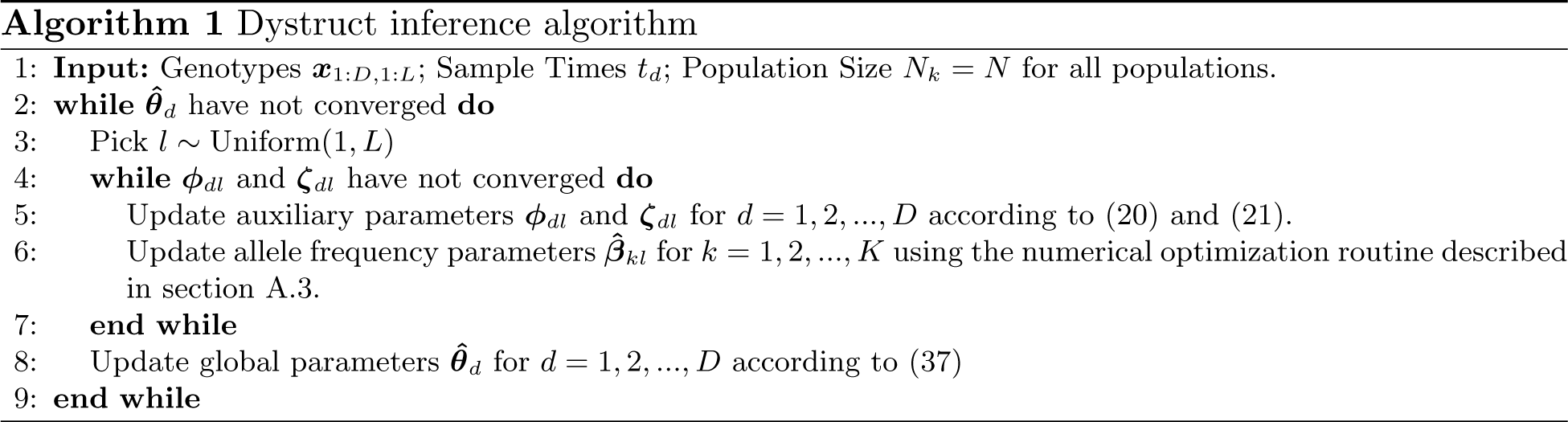

### A.5 Extensions to missing data

The above algorithm only holds for complete data. A small modification is required for missing data, where not every sample has an observed genotype at every locus. Rather than a single global step size *∊_t_*, we maintain a step size for every individual *∊_n__d_* where *n_d_* is the number of iterations for individual *d*. When a locus is subsampled, we only update global ancestry estimates for individuals with observed genotypes at that locus, and the step size for those individuals. We further replace the parameter *L* with *L_d_*, the number of loci for each observed in each individual.

